# Using Transcriptional Signatures to Find Cancer Drivers with LURE

**DOI:** 10.1101/727891

**Authors:** David Haan, Ruikang Tao, Verena Friedl, Ioannis Nikolaos Anastopoulos, Christopher K Wong, Alana S Weinstein, Joshua M Stuart

## Abstract

Cancer genome projects have produced multidimensional datasets on thousands of samples. Yet, depending on the tumor type, 5-50% of samples have no known driving event. We introduce a semi-supervised method called Learning UnRealized Events (LURE) that uses a progressive label learning framework and minimum spanning analysis to predict cancer drivers based on their altered samples sharing a gene expression signature with the samples of a known event. We demonstrate the utility of the method on the TCGA dataset for which it produced a high-confidence result relating 53 new to 18 known mutation events including alterations in the same gene, family, and pathway. We give examples of predicted drivers involved in TP53, telomere maintenance, and MAPK/RTK signaling pathways. LURE identifies connections between genes with no known prior relationship, some of which may offer clues for targeting specific forms of cancer. Code and Supplemental Material are available on the LURE website https://sysbiowiki.soe.ucsc.edu/lure.

## 1. Introduction

Cancer is a genetic disease caused by mutation and selection in somatic cells. Mutations in normal cells are usually repaired or result in apoptosis. Whereas in cancer cells mutations accumulate, leading to uncontrolled growth and tumorigenesis. There are two broadly defined types of mutations: drivers and passengers. Tumors contain around 2-5 driver mutations that cause and accelerate cancer, and about 10-200 passenger mutations which are accidental byproducts of thwarted DNA repair mechanisms.^1^ Driver mutations define some characteristics of the tumor and may offer therapeutic targets, yet identifying them amid the myriad of passengers remains a challenge.

The Cancer Genome Atlas (TCGA) is a publicly accessible dataset of cancer samples from the National Cancer Institute (NCI).^2^ TCGA catalogues mutations, mRNA, miRNA, DNA methylation, copy number variation, and protein expression data for roughly 11,000 patients across 33 cancer types.^2^ The identification of driver mutations played a key role in many TCGA analyses. For example, the TCGA study of papillary thyroid carcinoma identified two tumor subtypes characterized by different driver mutations. One subtype harbored mutations in BRAF and the other subtype in RAS genes such as KRAS, NRAS, or HRAS. This study identified at least one driver mutation in about 95% of the samples, leaving about 5% of the samples without known driver mutations.^3^ Here we present a semi-supervised pattern recognition tool, Learning UnRealized Events (LURE), that finds new drivers sharing molecular signatures with known drivers.

There are several existing computational tools that try to decipher driver from passenger mutations.^4^ EPoC uses network modeling of the transcriptional effects of copy number aberrations to identify driver mutations in glioblastoma (GBM).^5^ DriverNet employs a probabilistic model to locate driver mutations using transcriptional networks.^6^ These methods can predict novel drivers given a set of SNVs or copy number alterations and the corresponding mRNA gene expression data. In addition, there are methods that identify modules of driver genes based on mutual exclusivity in certain tumor types, such as CoMEt^7^ and MEMo, the latter of which incorporates prior knowledge such as pathway data into driver gene module discovery.^8^ In contrast, LURE uses mRNA data to identify mutations in “driver-unknown” samples with similar expression signatures to known drivers, thereby implicating a novel set of mutations as possible drivers.

Several studies have built gene expression signatures to identify samples with certain driver events. For example, studies have identified a TP53 gene expression signature as a reliable and independent predictor of disease outcome in breast cancer.^9,10^ In addition, in patients with epithelial ovarian cancer, a BRCAness gene expression signature is just as predictive of chemotherapy responsiveness and outcome as mutation status.^11^ While creating signatures as prognostic markers to guide treatment is important in a clinical setting, there has been little work using such gene expression signatures to find related mutational events. LURE identifies gene expression signatures across the 723 COSMIC cancer genes^12^ and then uses iterative semi-supervised learning to discover potentially related events.

## 2. Method

LURE associates events by finding similar molecular signatures among the samples in which the events occur. An event here is a particular type of alteration that affects a single gene, such as a focal deletion, a missense mutation, a truncating mutation, a gene fusion, and so forth. In this study, mRNA sequencing-based expression data is used to generate signatures, although other data choices are possible (e.g. microRNA expression or DNA methylation). LURE finds related events by training a classifier using the samples containing a known driver mutation (the “bait”). It then applies the classifier to find “target” samples defined as high-scoring samples lacking bait alterations. In classic machine-learning, the target samples would be considered false-positives. However, in this case, they offer an opportunity to find drivers because the comprehensive collection of TCGA mutation calls can be searched to find other events that significantly coincide with the target samples. Any such event (the “catch”) is associated with the starting bait and provides a new set of labels for retraining a more accurate classifier in subsequent rounds. Thus, the approach can be viewed as a specific kind of semi-supervised learning in which the possible labels are constrained to the event samples. LURE concomitantly identifies related events, expands the labels, and improves the classification accuracy using the iterative procedure described next.

LURE’s first step establishes a known driver mutation as the initial bait (Step 0; Figure 1A). Next, LURE trains a logistic regression classification model using gene expression as features and bait mutation status as the label to be predicted (Step 1). The method retains only those baits that yield accurate models determined by cross-validated area under the precision recall curve (PR AUC). LURE then uses the classification model to score each sample in the dataset (Figure 1A,B). Despite the inherent bias towards overfitting because the classifier is applied to samples used for training, some samples without the bait alteration may still receive a high classifier score, thus producing a sufficiently large set of target samples to identify and associate new catch events.

**Fig. 1.**
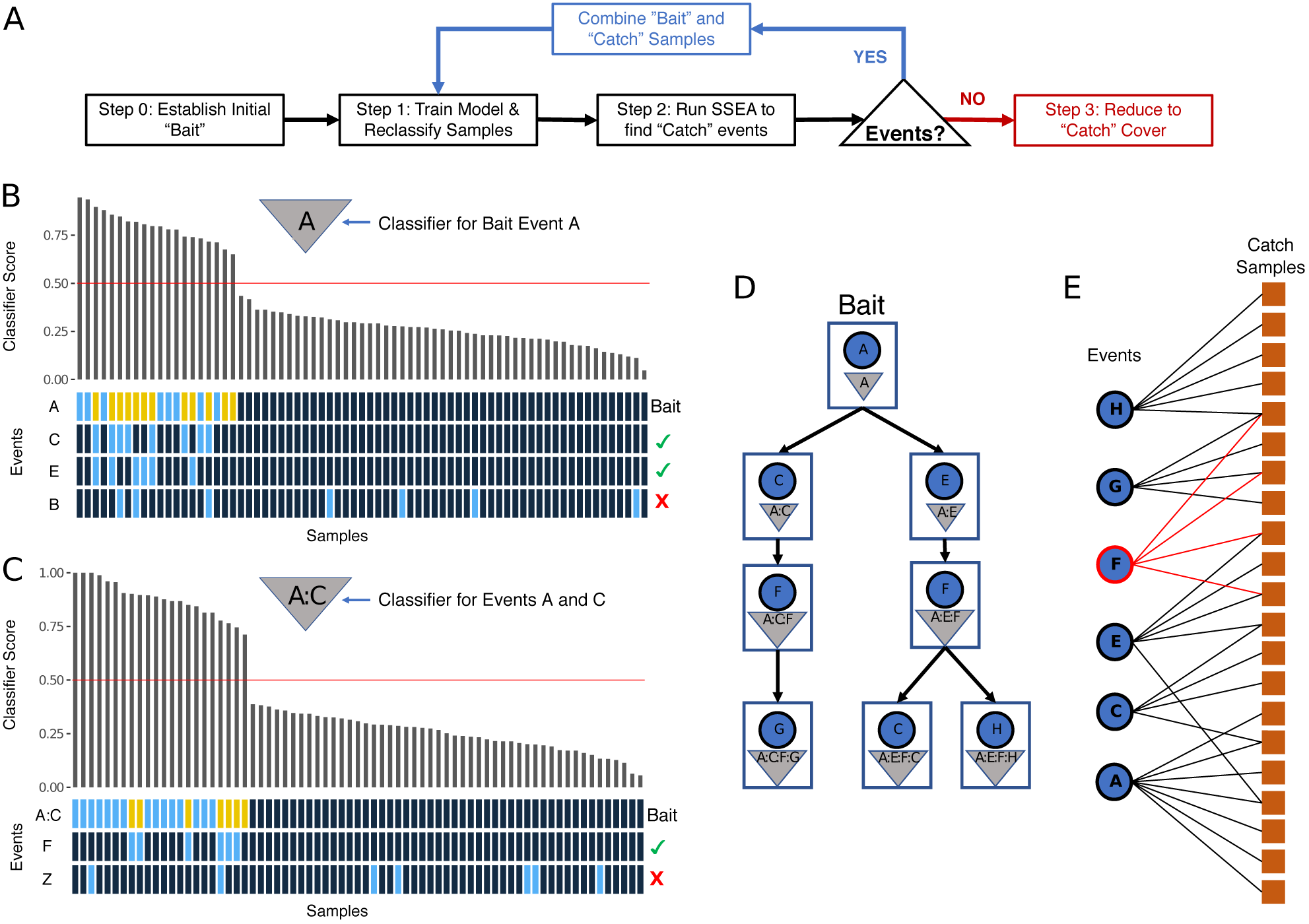
LURE Method Overview. **(A) LURE Method Flowchart. (B) LURE Graph of Bait Event A.** Scores (y-axis) assigned to samples (x-axis) by the logistic regression model trained to recognize samples harboring event A (triangle). Red line indicates 0.5 score cutoff for positive predictions. Presence of mutations for each sample are indicated below the barplot (blue, present; black, absent). Positive samples lacking alterations in the bait event define the set of “Target” samples (yellow). Events C and E are found by Sample Set Enrichment Analysis (SSEA) to be present in samples with high classifier scores (green check marks on right). **(C) LURE Graph of Combined Bait Event A:C.** Results from a classification model trained to recognize samples with either event A or C. New event F is found by SSEA to be associated with the A:C classifier. **(D) LURE Event Discovery Tree.** Each node represents an event (node label) discovered using the classification model (triangle) of its parent. The new classification model is trained using samples with the new event as well as any samples with events along the path to the root of the tree. The blue circles within each node represent the newest event added to the model. **(E) LURE Catch Cover.** A bipartite graph is constructed using all events from the Event Discovery Tree (left nodes) and all samples found to have scores greater than 0.5 with the final combined classification model as well as an alteration in one of the catch events (“Catch Samples”, right nodes). If an event is present in a sample, their nodes are connected in the graph. The set cover algorithm is run to obtain the minimal set of events that spans all samples. Event F illustrates an example of an event that would be removed by the algorithm because the samples with event F contain other events that explain the observed signature. The final event set is retained as the “Catch Cover.”

To leverage this new information, LURE runs a Sample Set Enrichment Analysis (SSEA) to test whether the samples of an event are associated with the target samples (Step 2, Figure 1A). For SSEA, the GSEAPreranked tool^13^ is used, which performs a Kolmogorov-Smirnov test to determine if samples with an event significantly sort toward the top of the list when all non-bait samples are ranked by their classification scores. LURE establishes a set of “Catch” events as those with significant SSEA association scores (p < 0.05, FDR < .25, number of events > 3). For each catch event, LURE then combines the positive samples for both bait and catch events into a new, intermediate bait event and trains a new classifier for these samples (Return to Step 1; Figure 1A). Cross-validation is run for the new classification model and the PR AUC results are compared to the initial classifier to ensure the model improves when including the new positive samples (Student’s t-test, *t* > 0). In addition, LURE tests the new classifier against a null model created by adding the same number of randomly chosen catch samples to the true positives and running cross-validation (Student’s t-test, *p* < 0.05). After establishing that the new event both improves the original classifier and significantly outperforms a random background distribution, the new classifier is rerun on all samples to search for the next set of catch events (Figure 1A,C). In this manner, LURE builds an “Event Discovery Tree” (EDT) by connecting catch events to the bait classifiers that discovered them, and recursively building new classifiers at each node(Figure 1D). The tree recursion stops when no further events are found by SSEA or the classifier performance no longer increases.

The above procedure can lead to the association of numerous events at every level of the EDT. By chance, mutations in passenger events can occur in the target samples, especially as a result of highly unstable genomes. Our task then is to distinguish such passengers from true, but as-yet unknown, drivers. To do this, LURE identifies a set of events, where each event occurs in at least some target samples that contain no other event. Intuitively, these events may be the sole explanation for driving the signature in the target samples. To make this concrete, consider the hypothetical scenario in which two events — X and Y — have been included in the EDT. Further, imagine that several samples have mutations only in event X. In contrast, no such samples exist for event Y; i.e. Y only occurs in samples that have at least one other event from the EDT. LURE assumes that X is more likely to be a driver than Y because X is the only event that can explain the presence of the signature in several samples.

Thus, in the last step (Step 3; Figure 1A), LURE computes a minimal set of events in the catch that best span the catch samples. To this end, LURE builds a final classification model from the union of all events in the EDT and uses this model to identify a final set of samples scoring higher than 0.5. It then takes all events in the EDT that span these high-scoring samples (Figure 1E) and runs the Vazirani approximation to the set cover algorithm^14^ as implemented in the RcppGreedySetCover R Package. The set cover algorithm identifies the minimum set of events, the “Catch Cover,” that collectively occur in all catch samples, removing events with completely overlapping samples in order to enrich for events that are uniquely responsible for the expression signature.

More details about LURE and its implementation are available in the Supplemental Methods section on the LURE website https://sysbiowiki.soe.ucsc.edu/lure.

## 3. Results

### 3.1 Datasets

All data were obtained from the TCGA Pan-Cancer Atlas collection.^2^ These data included batch-corrected and normalized gene expression data, as well as single nucleotide variant (SNV) calls determined from the TCGA’s MC3 consensus analysis, and gene-associated copy number data from GISTIC2.

### 3.2 Positive Controls for LURE

Isocitrate Dehydrogenase 1 (IDH1) is one of three IDH enzymes, which when mutated causes hypermethylation and subsequent altered gene expression in gliomas.^15^ In order to test LURE’s ability to discover known “catch” events, we created a test set of bait and catch events using IDH1 missense mutations in the TCGA Lower Grade Glioma (LGG) sample set. Of the 210 LGG samples with an IDH1 missense mutation, we created an initial bait with 150 samples and three sets of 20 samples as potential catch events. LURE identified all three of the held-out events in the Catch Cover, as well as the IDH2 missense mutation event. IDH2 is also one of the three IDH isozymes, and a mutation in IDH2 has the same oncogenic effect as an IDH1 mutation^16^(Supp. Figure 1A). We also tested a similar method called REVEALER on this positive control set. We found that REVEALER was unable to identify the hold-out samples (Supp. Figure 2).

**Fig. 2.**
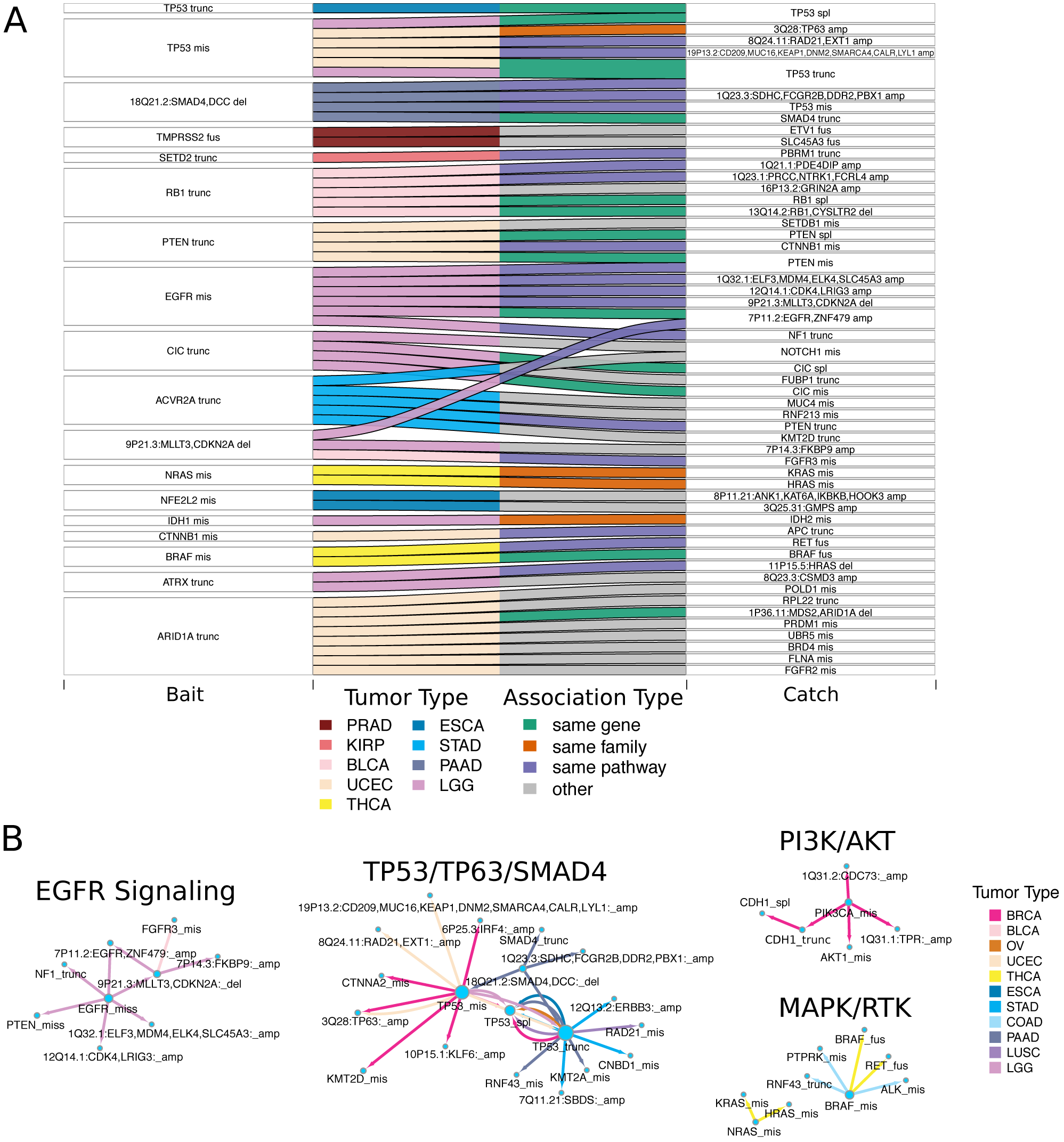
LURE Pan-Cancer Results. (A) High Confidence Bait-Catch Associations. Sankey diagram shows the high confidence bait-catch associations for the 18 baits with a final classifier PR AUC *>* 0.8. Bait gene and mutation type are shown on the left; catch gene and mutation type on the right. Each horizontal flow connection represents an association between the bait and catch event found by LURE. The left half of each flow bar is colored by tumor type in which the association was found. The right half of each flow bar is colored by the association type. **(B) LURE “Event Net” Showing Selected Pathway Associations.** Cytoscape^21^ visualization of a subset of LURE associations in the TCGA Pan-Cancer dataset grouped by pathway. Each node represents an event. Directed edges represent an association and the direction of the LURE discovery (bait to catch). The color of each edge represents the tumor type in which the association was found. Refer to Supp. Figure 8 for full Cytoscape visualization of all Pan-Cancer results.

Splicing Factor 3b Subunit 1 (SF3B1) is a well-known splicing factor which is recurrently mutated in many tumor types, including Uveal Melanoma (UVM).^17^ Missense mutations in SF3B1 lead to aberrant splicing and a unique gene expression signature.^18^ We used SF3B1-mutated tumors in the TCGA UVM dataset as a positive control. Of the 80 UVM samples, 18 samples have missense mutations in SF3B1. We created an initial bait using 8 SF3B1-mutated samples and left out two sets of 5 SF3B1-mutated samples for discovery. LURE re-discovered both withheld sets correctly, collecting all of the SF3B1 missense events in the Catch Cover (Supp. Figure 1B).

### 3.3 LURE on the TCGA Pan-Cancer Dataset

In order to look for novel associations among genes already implicated in cancer, we ran LURE across all tumor types in the TCGA Pan-Cancer Atlas dataset, restricting both baits and catches to mutation events in the 723 COSMIC genes.^2,12^ We created bait events for missense mutations, truncating mutations, homozygous focal point copy number deletions, splice site mutations, and gene fusions. We only considered mutations that occurred in at least 10 samples for a given tumor type. We created tumor type-specific classification models to avoid any confounding effects due to tissue-specific expression and mutation patterns. By creating baits for different alteration types in the same gene, as opposed to one bait for any alteration in a gene, we were able to identify associations between different alteration types within the same gene as well as infer the functional impact of alterations. For example, *α*-thalassemia mental retardation X-linked (ATRX), a gene recurrently mutated in LGG, only has an oncogenic effect with a loss-of-function mutation — either a truncating mutation or copy number loss — whereas a missense mutation may not have an oncogenic effect.^16^

We trained logistic regression models on the resulting 3,053 bait/tumor type combinations. We also tested both random forest and neural network models and found the logistic regression model to consistently score higher for the majority of mutations (Supp. Figure 3). Due to the imbalanced nature of the data, precision and recall, rather than overall accuracy, was used to measure classifier accuracy to emphasize the ability of a classifier to detect bait samples that are often highly underrepresented.^19^

**Fig. 3.**
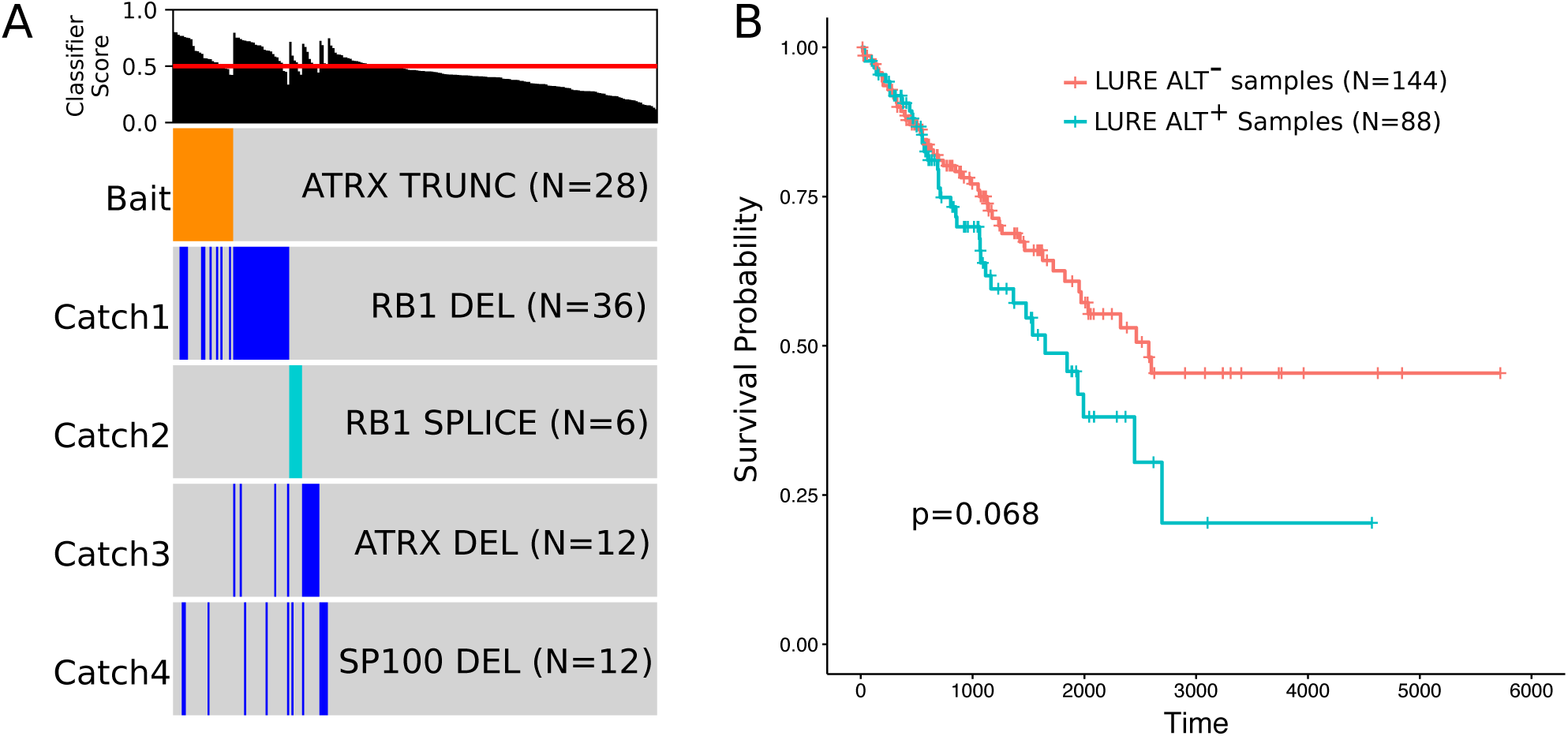
LURE results for the Alternative Lengthening of Telomeres (ALT) in Sarcomas. (A) LURE ALT Graph. Using a known driver for ALT (ATRX truncating mutations) as bait in Sarcomas (SARC), LURE finds four catch events: copy number deletions of ATRX, RB1, and SP100, as well as RB1 splice site mutations. **(B) LURE ALT Survival Plot** Kaplan-Meier survival plot dividing SARC samples by ALT classification using the final Catch Cover classifier from LURE. LURE ALT^*-*^ samples have classification scores *<* 0.5, ALT^+^ samples *≥* 0.5.

In order to limit the number of putative false positive results, we restricted the number of bait classifiers by considering only those with PR AUC > 0.5, precision > 0.4, and recall > 0.75 (Supp. Figure 4). Since the objective of LURE is to identify and reclassify target samples, we were more lenient with the precision cutoff and placed more restriction on the recall cutoff. Among the bait classifiers passing these thresholds, the most common bait gene across all tumor types was againTP53, and the tumor types with the highest number of passing bait classifiers were Lower Grade Glioma (LGG), Thyroid Carcinoma (THCA), and Prostate Adenocarcinoma (PRAD) (Supp. Figures 5,6).

**Fig. 4.**
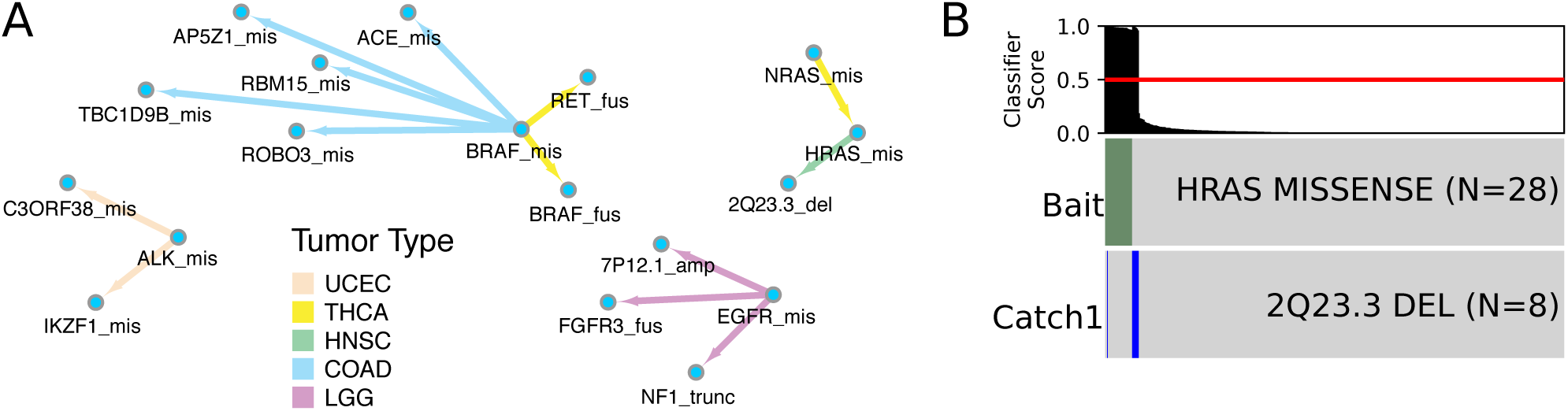
LURE RAS/MAPK Signaling Pathway Analysis. **(A) LURE “Event Net” Showing Associations with MAPK/RTK signaling pathway genes.** Each node represents an event. Directed edges represent associations and the direction of the LURE discovery (bait to catch). The color of each edge represents the tumor type in which the association was found. **(B) LURE Graph of HRAS in HNSC.** Using HRAS missense mutations as bait in Head and Neck Squamous Cell Carcinomas (HNSC), LURE found a deletion catch event in the 2q23.3 locus.

After filtering the models, we ran LURE with the 81 remaining classifiers as bait (Supp. Figures 5,6), using missense mutations, truncating mutations, splice site mutations, gene fusions, focal point copy number amplifications, and homozygous deletions of COSMIC genes as possible catches. LURE found significant bait-catch associations for 35 of the 81 baits tested. Adding catches to each initial bait event increased the classifier PR AUC by various amounts across the different baits (Supp. Figure 7A). Tumor type was evenly distributed among the baits and SNVs dominated the bait mutation type (Supp. Figure 7D). The most common gene among the 35 baits was again TP53 (Supp. Figure 7E).

Among the high-confidence results with a final classifier PR AUC *>* 0.8, 14 of 59 individual bait-catch associations were between events involving the same gene, such as TP53 truncating, splice site, and missense mutations, in various tumor types (Figure 2A). There were four associations within the same gene families, e.g. IDH1/2 or the RAS protein family. In addition, we identified gene fusion event partners in BRAF and RET that associated with a BRAF missense mutation in THCA. For 20 of the 59 high confidence bait-catch associations, both genes were members of the same signaling pathway (excluding pathway gene sets with *>* 1000 genes).^20^

When all pan-cancer results are considered together, the resulting bait-catch association network, or “Event Net,” reveals some pathway-oriented findings which cross multiple tumor types (Supp. Figure 8). In particular, four canonical pathway-oriented findings emerge: EGFR signaling, TP53/TP63/SMAD4, PI3K, and MAPK/RKT (Figure 2B). LURE identified interesting associations for PTEN, in particular between PTEN and CTNNB1, a connection supported by recent research which suggests PTEN plays a role in regulating the subcellular localization of *β*-catenin.^22^ Another striking LURE association is between PTEN and EGFR, which is consistent with recent findings suggesting that PTEN regulates EGFR signaling.^23^ These LURE associations for PTEN reveal crosstalk between pathways and provide further evidence that alterations in PTEN influence EGFR signaling and *β*-catenin signaling.

### 3.4 LURE Finds New Drivers of the Alternative Lengthening of Telomeres Pathway in Sarcomas

Tumor cells must employ Telomere Maintenance Mechanisms (TMMs) to extend their telomeres in order to multiply rapidly and avoid senescence.^24^ To date, there are two known mechanisms tumor cells use to avoid telomere erosion: the overexpression of telomerase, an enzyme with the ability to extend telomeres, and the Alternative Lengthening of Telomeres (ALT) pathway. The vast majority of tumors overexpress telomerase in some way, whereas a small portion (10-15%) use ALT.^25^ ALT-positive (ALT^+^) samples lengthen telomeres through homologous recombination, mediated by loss-of-function mutations in the ATRX and DAXX genes.^26^ Approximately 80% of tumors with ALT harbor mutations in ATRX or DAXX,^27^ leaving 20% with no known driver. Using LURE, we sought to identify new driver mutations of the ALT pathway using gene expression signatures of samples harboring ATRX loss-of-function mutations. Sarcomas and Lower Grade Gliomas (LGG) have the highest prevalence of ALT^+^ samples, and ATRX is recurrently mutated in these diseases. We therefore chose these tumor types in which to search for new drivers of the ALT pathway.^26^ We restricted our gene set to a manually-curated set of genes associated with telomere maintenance derived from the TelNet database^28^ and focused on looking for associations with ALT. Since TP53 is commonly mutated in ALT^+^ samples and TP53 mutations are not known to be sufficient to cause ALT,^29^ we excluded TP53 events from the possible catches to identify novel ALT drivers.

Using ATRX truncating mutations as bait, LURE identified four associated mutations in sarcomas (Figure 3A). While we would expect ATRX truncating mutations to associate with an ATRX copy number deletion,^30^ the deletions found in RB1 and SP100 are novel. We suggest the expression signature LURE identified in this analysis is classifying ALT^+^ TMM samples, and the associated alterations are possibly driving the TMM. Previous work has associated RB1 alterations with long telomeres in the absence of TERT mutations and ATRX inactivation.^31^ In addition, mouse models have revealed that the knock-out of Rb-family proteins causes elongated telomeres.^32^ LURE also identified SP100 deletions as an ALT driver, and while SP100 deletions have not been directly reported to be involved in ALT, overexpression of SP100 in ALT^+^ cell lines has resulted in suppression of ALT characteristics.^33^ We therefore suggest that an SP100 deletion may lead to unhindered ALT TMM activity. To further investigate the subset of LURE-classified ALT^+^ samples, we performed a survival analysis and found that the ALT^+^ samples show a worse prognosis (*p* = 0.068) (Figure 3B),

In LGG, again using ATRX truncating mutations as bait, LURE found similar results identifying associations with other ATRX alterations such as ATRX deletions, splice site mutations, and missense mutations (Supp. Figure 9). While ATRX missense mutations are not generally thought to be ALT drivers, we found that ATRX missense mutations were in fact associated with truncating mutations, suggesting a loss of function role for some of these missense mutations.^30^ Together, these findings implicate new single and combinations of driver mutations required for the initiation of the ALT telomere maintenance mechanism and could prove to be therapeutic targets.

### 3.5 LURE Identifies Associations within the MAPK/RTK Signaling Pathway

Oncogenic mutations of the HRAS, NRAS, or KRAS genes are frequently found in human tumors, altering the control of cellular proliferation, differentiation, and survival. Oncogenic mutations in a number of other upstream or downstream components of the MAPK/RTK signaling pathway, including membrane receptor tyrosine kinases (RTKs) and cytosolic kinases, have been recently detected in a variety of cancer types.^35^ Oncogenic RAS mutations and other mutation events within the MAPK/RTK signaling pathway are often mutually exclusive, indicating that the deregulation of Ras-dependent signaling is essential for tumorigenesis.^35^ Previous studies have shown that tumor samples harboring Ras protein mutations have a unique gene expression signature, and Ras-dependent samples can be more accurately defined by this signature than by mutation status alone.^36^ Building on this knowledge, we were able to use LURE not only to train an accurate Ras-dependent classifier, as was done in Way et al,^37^ but also to identify new alterations which may be activating the MAPK/RTK signaling pathway in samples without a Ras protein mutation. We ran LURE using genes known to be involved in the MAPK/RTK signaling pathway^38^ as baits. We considered baits with the following alterations: missense mutations, truncating mutations, homozygous focal point copy number deletions, splice site mutations, and gene fusions. We did not restrict our catch set to COSMIC genes as we were looking for genes not previously implicated in cancer. We ran LURE for the 23 bait event classifiers which scored greater than 0.5. The resulting Event Net revealed known as well as novel associations (Figure 4A). One interesting association found by LURE in Head and Neck Squamous Cell Carcinomas (HNSC) is between HRAS missense mutations and focal deletions of the 2q23.3 locus (Figure 4B). The samples were almost mutually exclusive for these events, with only one sample having both a 2q23.3 deletion and an HRAS alteration and no samples having alterations in either KRAS or NRAS. Among the 61 genes in the 2q23.3 locus is CHST11, which has been shown to regulate MAPK/RTK pathway activity in hepatocellular carcinoma.^39^ We suggest that in the absence of an HRAS mutation, MAPK signaling may be activated by a deletion of the 2q23.3 locus in HNSC.

## 4. Discussion

We described the Learning UnRealized Events (LURE) method that leverages the availability of multiomic cancer datasets to identify driver events using signatures constructed from other feature data (in this study, mRNA expression). LURE is related to semi-supervised machine learning approaches,^40^ but has a slight shift in emphasis. Rather than focusing on labeling unknown samples to optimize performance, LURE instead uses the mutual compatibility of label sets to form a coherent classifier as an indicator that the label sets themselves are related. Thus, the accuracy of the classifier is only important to the extent that it produces a signature that can detect strong associations. LURE takes advantage of the “dark matter” (false-positives in gene expression-based classifiers) to search for associated events.

LURE outperformed REVEALER, another signature-based method, on the IDH1 positive control test. REVEALER identifies combinations of genomic alterations correlated with functional phenotypes, such as the activation or gene dependency of oncogenic pathways or sensitivity to a drug treatment.^41^ While the concept of REVEALER is similar to LURE, LURE provides several notable advantages that may lead its better performance. First, at every iteration of the method, LURE produces a new classification model that is more accurate than the model from the previous iteration. LURE does this by expanding the training set to include samples with alterations in the new event, which in turn updates the signature with potentially new features to aid classification. Second, REVEALER’s predictions rely on mutually exclusive relationships between new events, and its results are thus limited by the accuracy of mutation calls. By allowing some overlap between predicted events, LURE can account for possible mutation call errors and identify modules containing co-mutated events.

Some limitations of this study could be addressed to expand the set of predicted drivers. First, we used a highly conservative set of parameters to analyze the TCGA dataset in order to control the set of false positive results and to highlight the most confident connections. These include the minimum allowed accuracy of the bait classifiers, the cutoff for the SSEA tests, the allowed overlap between the samples of the events, and restricting the considered catch to COSMIC genes. To obtain different candidates, one could run a less stringent LURE analysis to produce many more catch events that could be further filtered using criteria such as the recurrence of the same catch found in multiple tumor types. Second, we only considered gene expression data as features for building signatures. Yet multiple other data types are available, such as DNA methylation, to increase the ability to find associations among events. Third, the use of SSEA restricts the consideration of catch events to those occurring in at least four target samples in order to pass the multi-test FDR cutoff. However, it might be worthwhile to add events to the catch even if they cover only one additional target sample. Methods that consider sets of mutually exclusive events simultaneously might offer some advantage.

Using signatures to associate events will extend the list of known drivers that may ultimately identify targets for precision medicine. LURE identified an intriguing set of new candidate drivers based on the Pan-Cancer Atlas dataset. Most of the found associations were between events of the same gene (e.g. TP53), gene family (e.g. IDH, RTK), or biological pathway (e.g. PI3K). In addition to associating known events across the Pan-Cancer dataset, LURE identified many events of unknown significance to known pathways such as putative drivers of telomere maintenance and MAPK/RTK signaling. The collection of these events provide possible actionable clues for cancer patients. For example, LURE found that deletions in 2q23.3 in head-and-neck cancers are strongly associated with RTK signaling. Whether such a focal deletion could be used as a novel biomarker for treating patients with an RTK inhibitor, such as gefitinib for EGFR, remains to be seen.

## Notes

https://sysbiowiki.soe.ucsc.edu/lure

